# Inductive inference of novel protein-molecule interactions using Heterogeneous Graph Transformer (HGT) AutoEncoder

**DOI:** 10.1101/2021.12.20.472904

**Authors:** Alberto Arrigoni

## Abstract

Protein-molecule interactions are promoted by the physicochemical characteristics of the actors involved, but structural information alone does not capture expression patterns, localization and pharmacokinetics. In this work we propose an integrative strategy for protein-molecule interaction discovery that combines different layers of information through the use of convolutional operators on graph, and frame the problem as missing link prediction task on an heterogeneous graph constituted by three node types: 1) molecules 2) proteins 3) diseases. Physicochemical information of the actors are encoded using shallow embedding techniques (SeqVec, Mol2Vec, Doc2Vec respectively) and are supplied as feature vectors to a Graph AutoEncoer (GAE) that uses a Heterogeneous Graph Transformer (HGT) in the encoder module. We show in this work that HGT Autoencoder can be used to accurately recapitulate the protein-molecule interactions set and propose novel relationships in inductive settings that are grounded in biological and functional information extracted from the graph.

## I. Introduction and previous work

**I**N ORDER TO TO interact with and influence the expression of a target protein, a chemical compound needs to 1) access the same bodily (e.g. the blood-brain barrier) and cellular compartment of the target [4] 2) being absorbed and degraded with a kinetic that is compatible with the metabolism of the target 3) present a chemical structure that is suitable for binding to some active site on the target protein.

Much of the literature in this domain focuses on requirement 3) [16], whereas the other factors are also essential to exert any kind of biological function. Hence, focusing solely on the chemical properties of the actors results in the selection of molecules which fall short on the promise to modulate the functionality of e.g. a target enzyme, although they may perform well in *in vitro* assays. Moreover, when dealing with a complex pathway of interacting proteins, it is not clear how to select a suitable target: the intrinsic collaborative fashion through which genes and proteins interact may lead to the emergence of ‘salvage routes’[3], which render the modulation of a single component ineffective. This notion is inherently stored in the vast wealth of knowledge built upon the known interactions between molecules and proteins, and in turn proteins with other proteins. We must add to this information the existing *mechanistic* knowledge regarding the role that a protein plays in the context of a particular disease.

### A. Current work contributions

When predicting the potential for a molecule to interact with a certain protein (or pathway) we can frame the problem as ‘missing link prediction’ task of an heterogeneous graph constituted by three entities: 1) molecules 2) proteins 3) diseases (fig.I-A).

**Fig. 1.**
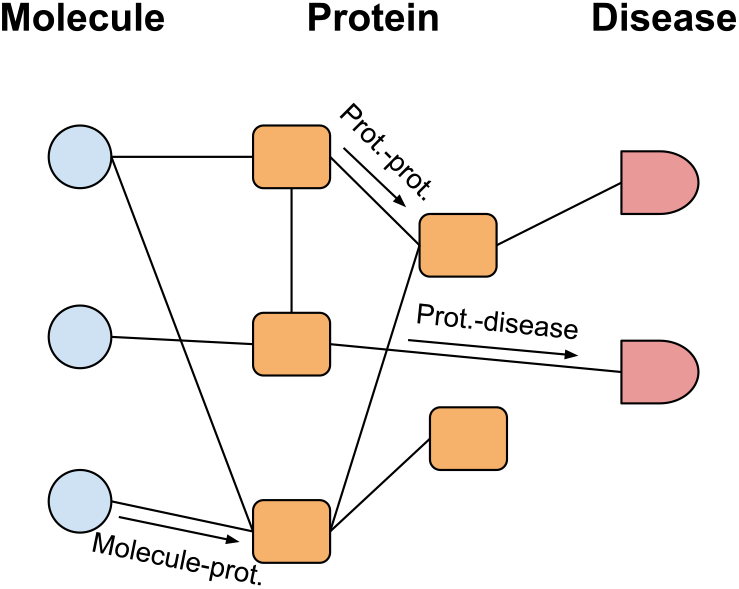
Structure of the heterogeneous graph used in this work. The relationships listed are used for message passing

In order to integrate physicochemical data with interactomic information stored in the graph we calculate shallow embeddings for each entity (using respectively MOL2VEC, SeqVec and Doc2Vec) and feed these vectors to the downstream model.

We wish to address the requirements for inference of novel protein-molecule interactions by using a modified version of a Graph Autoencoder (GAE) which handles heterogeneous data and performs inductive inference by the means of an encoder structure that uses a stack of Heterogeneous Graph Transformer (HGT) convolutional layers. We show that HGT GAE performs consistently better than popular alternatives (SAGE [7], GAT [18]), and it can be used for inductive inference on novel molecules/proteins.

## II. Materials and methods

### A. Datasets

We derived the drug-target interaction network from the publicly available datasets availble on SNAP [21]. The selected network aggregates high-throughput experimental data, manually curated datasets, and the results of several prediction methods into a single global network of chemical-gene interactions. The number of drug nodes is 1774, the protein nodes are 17984 and the number of edges between them is 131034. The protein-protein network was downloaded from the SNAP data repo as well. It includes direct (physical) proteinprotein interactions, as well as indirect (functional) associations between human proteins. Nodes represent proteins and edges represent associations between them. Disease description, disease class and protein associations were taken from DisGeNET [13].

### B. Node features embedding per entity type

Each one of the entity types represented in the graph bears useful information in addition to the graph context it resides in. For 1) proteins (*p*_*i*_) is the aminoacid sequence which is related to the overall secondary and tertiary structure, as well as active sites architecture, for 2) molecules (*m*_*i*_) is the chemical properties, while for 3) diseases (*d*_*i*_) is a general textual description related to the characteristics of the disease and often its bodily localization (e.g. colorectal cancer).

Embeddings of this information are generated and then used as feature vectors in the message passing procedure performed by the encoder (see fig.II-C1A).

**Fig. 2.**
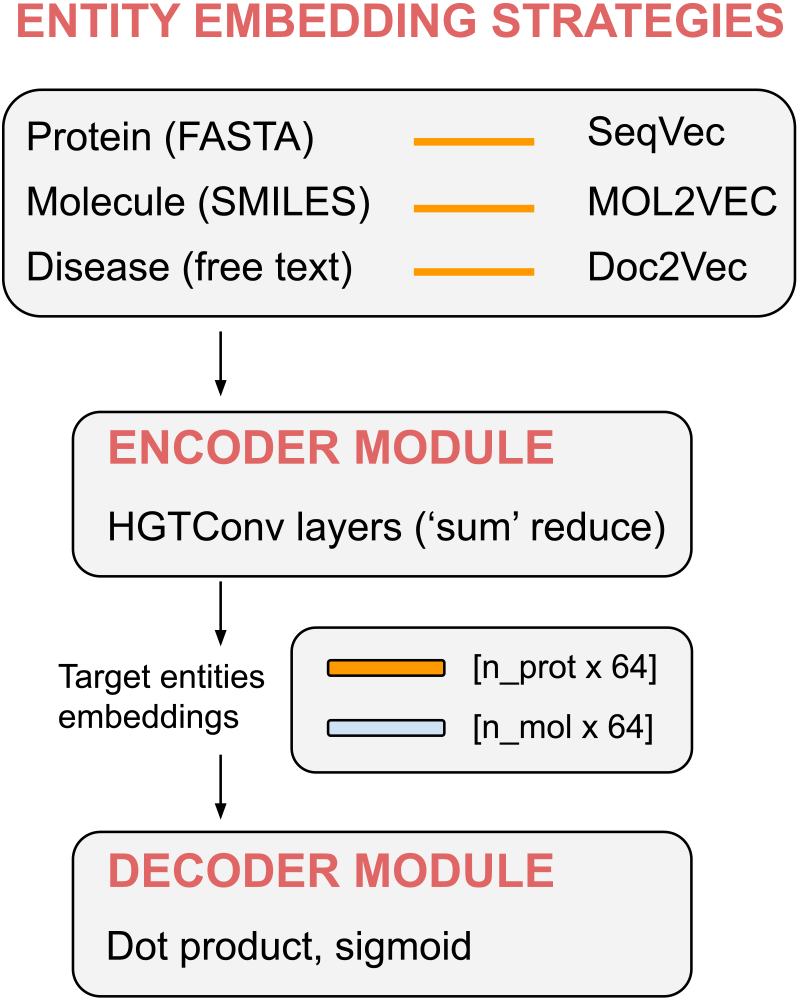
A) Embedding strategies. SeqVec[8] was used on FASTA protein sequences, MOL2VEC [10] on SMILES molecule sequences and Doc2Vec [11] on the textual representation of disease descriptions B) Model architecture

1. SeqVec[8] was used on FASTA protein sequences to produce vector embedding *p*_*i*_ ∈ ℝ^1024^. This approach exploits the deep bi-directional model ELMo taken from natural language processing (NLP) to generate embedddings that capture the biophysical and biochemical properties of protein sequences from unlabeled big data (UniRef50).
2. MOL2VEC [10] on SMILES molecular sequences to produce vector embedding *m*_*i*_ ∈ ℝ^300^. MOL2VEC is an unsupervised machine learning approach that learns vector representations of molecular substructures that point in similar directions for chemically related substructures.
3. Doc2Vec [11] on the textual representation of disease descriptions to generate vector embedding *d*_*i*_ ∈ ℝ^5^). Disease description, disease class and protein associations were taken from DisGeNET [13].

### C. Model architecture

The model follows the architecture of a Graph AutoEncoder (GAE), which is used in different contexts to predict missing associations between nodes [2], [14].

As the graph built for this work is heterogeneous, the message-passing aggregations are combined across different node types through a reduce operation. The encoder is composed of a series of convolutional operators (Heterogeneous Graph Transformer [HGT]) which produce embeddings of the entities between which we want to predict missing associations (molecules, proteins). The decoder performs the dot product of embedding vectors whose output is fed to a cross-entropy loss function that evaluates the reconstruction capabilities of the model end-to-end. As in classical GAEs, the goal is to reconstruct the original adjacency matrix (*A*) assuming missing links refer to unknown associations between molecules and proteins.

#### Definition 1. HeterogeneousGraph

A heterogeneous graph is defined as a directed graph *G* = (*V, E, A, R*) where each node *v ∈ V* and each edge *e ∈ E* are associated with their type mapping function *τ* (*v*) : *V* → *A* and *φ*(*e*) : *E* →*R*, respectively.

##### GraphNeuralNetwork

(general framework): we can generalize the convolutional operator in a graph context by expressing it as a neighborhood aggregation or message passing scheme. With 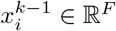 denoting node features of node *i* in layer (*k* ™ 1) and *e*_*j,i*_ ∈ ℝ^*D*^ denoting edge features from node *j* to *i*, message passing graph neural networks can be described as

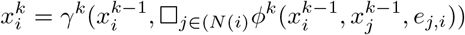

where □ denoted a differentiable, permutation invariant function, an **aggregation** function e.g. sum, mean or max, and *γ* and *phi* denote differentiable functions such as MLPs (Multi Layer Perceptrons). *φ* represents an **extraction** operation, which blends information originating from the neighbours with the target node’s features.

##### 1) Encoder

The encoder structure is modeled after the Heterogeneous Graph Transformer (HGT) [9] architecture. HGT builds node representations for proteins, molecules and diseases by performing heterogeneous **mutual attention** and heterogeneous **message passing**. Given a sampled heterogeneous sub-graph, HGT extracts all linked node pairs, where target node *t* is linked by source node *s* via edge *e*, identifying meta relation (*t, e, s*).

Heterogeneous mutual attention uses a distinct set of projection weights for each meta relation to calculate the attention matrix for *h* heads (**Attention**). Parallel to the calculation of mutual attention, heterogeneous message passing uses a typespecific projection matrix and concatenates all *h* messages for each node pair (**Message**).

The next step consists of a target-specific aggregation step that uses the attention vector as the weight to average the corresponding messages from the source nodes and get the updated vector representation of node *t, H*:

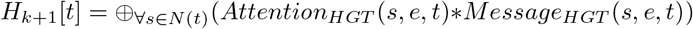

where *N* is a function that lists the neighboring nodes of *t* to aggregate information to the target node from all its neighbors.

The output of the encoder are two tensor both of shape (*n, e*), where *n* is number of proteins/molecules in the batch and *e* is the embedding tensor length.

### D. Decoder and loss function

The decoder uses the dot product between protein and molecule representations followed by a sigmoid activation function to generate a scalar output that is fed to the loss function.

Model parameters were optimized using the cross-entropy loss:

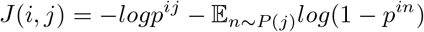

where (*i, j*) represents an observed edge, while (*i, n*) is a random edge obtained through negative sampling, as in [12].

## III. Results

### A. Benchmarking against related methods

In order to comparatively assess the results obtained on the graph dataset using different methods the Brier score was used. This metric measures the accuracy of probabilistic predictions, and it is used instead of e.g. AUC when a *calibrated* measure of error is needed. A calibrated model is needed for a binary classifier when not only the ranking of the outcomes is required, but also their relative distance in terms of importance. The Brier score is formulated as:

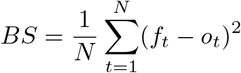

in which *f*_*t*_ is the probability that was forecast, *o*_*t*_ the actual outcome of the event at instance *t* (0 if it does not happen 1 if it happens) and N is the number of forecasting instances. Therefore, the lower the Brier score is for a set of predictions, the better the predictions are calibrated.

The HGT encoder was benchmarked against related methods that could be used to predict novel graph associations: a baseline multi-layer perceptron model (MLP), SAGE [7] and GAT[18]. Both SAGE and GAT are inductive methods that can be similarly used for link prediction on heterogeneous graphs. Results are shown in the table below:

**TABLE 1.** HGT performance (Brier score) compared with MLP, SAGE and GAT (listed performance is representative of 10 runs each)

| MLP (baseline) | SAGE encoder | GAT encoder | HGT encoder |
| --- | --- | --- | --- |
| 0.097 ± 0.005 | 0.048 ± 0.06 | 0.055 ± 0.063 | **0.032 ± 0.015** |

### B. Inferred relationships: a qualitative analysis

What follows are instances of downstream analyses that exemplify the way in which link prediction on graph based on prior *embedded* information can be used to uncover (or support) novel biological associations.

One way the system could be queried is by focusing on strategies to repurpose existing drugs for which toxicological and pharmacokinetic data are abundant. Paracetamol (also known as acetaminophen) [1] is a medication used to treat fever and mild to moderate pain. Being a drug of choice for reducing fever, it exerts its effects by inhibiting the cyclooxygenase and altering the actions of its metabolite AM404[5]. Among the top gene hits reported by the model for paracetamol we find MSH2, which is a component of the post-replicative DNA mismatch repair system. Evidence of the existing modulatory interaction between paracetamol and MSH2 is provided by a recent paper (2021)[19], confirming the validity of the predicted interaction. Similarly, an interaction between paracetamol and TYRP2 is predicted with high confidence and reported by http://genome.cse.ucsc.edu/cgi-bin/hgGene?org=Human&hgg_chrom=none&hgg_type=knownGene&hgg_gene=uc001vlv.6, although not included in the original dataset used for training and testing the model.

Most notably, one top predicted interaction of paracetamol, MAPK1, may shed light on liver toxicity effects found in patients following drug overdosage[15]. Paracetamol mediates hepatotoxicity by modulating the JNK signaling pathway, and it is the MAPK module itself that activates c-JUN N-terminal kinase by sequential protein phosphorylation.

To further exemplify the use of graph-derived insights to uncover novel applications and drug repurposing, we report that one of the top hits for Apixaban (a well known anticoagulant) is NR4A2. This protein is involved with others in the amplification of thromboinflammatory endothelial responses to the viral RNA analogue poly(I:C) [17], which in turn promotes the localized activation of platelets and the blood coagulation mechanism. This is also true for some of the inflammatory consequences found in Covid-19 patients, for which Apixaban is used as anticoagulation therapy [20].

Interestingly, the predicted interaction between apixaban and PDK1 was also recently reported by a different computational screening for the identification of schistosomicidal agents [6].

## IV. CONCLUSIONS AND FUTURE WORK

In this work we show how molecule-protein interactions can be inferred by integrating physicochemical and interactomic data through the use of shallow embedding methods and HGT AutoEncoders.

It is to be noted that the information that can be integrated in this graph model is greater than what portrayed in this proof-of-concept study: enzymes active sites location and description, expression level profiles, genomic coordinates and the presence of transcription factor binding motifs are only a few examples of elements that would add predictive and explanatory power to the model.

## Notes

### Competing Interest Statement

The authors have declared no competing interest.

